# Novel insights into the taxonomic diversity and molecular mechanisms of bacterial Mn(III) reduction

**DOI:** 10.1101/695007

**Authors:** Nadia Szeinbaum, Brook L. Nunn, Amanda R. Cavazos, Sean A. Crowe, Frank J. Stewart, Thomas J. DiChristina, Christopher T. Reinhard, Jennifer B. Glass

## Abstract

Soluble ligand-bound Mn(III) can support anaerobic microbial respiration in diverse aquatic environments. Thus far, Mn(III) reduction has only been associated with certain *Gammaproteobacteria*. Here, we characterized microbial communities enriched from Mn-replete sediments of Lake Matano, Indonesia. Our results provide the first evidence for biological reduction of soluble Mn(III) outside the *Gammaproteobacteria*. Metagenome assembly and binning revealed a novel betaproteobacterium, which we designate “*Candidatus* Dechloromonas occultata.” This organism dominated the enrichment and expressed a porin-cytochrome c complex typically associated with iron-oxidizing *Betaproteobacteria* and a novel cytochrome c-rich protein cluster (Occ), including an undecaheme putatively involved in extracellular electron transfer. This *occ* gene cluster was also detected in diverse aquatic bacteria, including uncultivated *Betaproteobacteria* from the deep subsurface. These observations provide new insight into the taxonomic and functional diversity of microbially-driven Mn(III) reduction in natural environments.

**Originality-Significance Statement:** Recent observations suggest that Mn(III)-ligand complexes are geochemically important in diverse aquatic environments. Thus far, microbially-driven Mn(III) reduction has only been associated with *Gammaproteobacteria* encoding three-component outer-membrane porin-cytochrome c conduits. Here, we demonstrate that *Betaproteobacteria* dominate in abundance and with respect to protein expression during biologically-mediated Mn(III) reduction in an enrichment culture from an anoxic lacustrine system. Using metaproteomics, we detect for the first time that *Betaproteobacteria* express a two-component porin-cytochrome c conduit, and an uncharacterized extracellular undecaheme (11-heme) c-type cytochrome. Although this is the first definitive report of an undecaheme within the *Betaproteobacteria*, we find evidence that they are widespread in uncultivated strains. These results widen the phylogenetic diversity of Mn(III)-reducing bacteria, and provide new insights into potential molecular mechanisms for soluble Mn(III) reduction

## Introduction

Manganese(III) is a strong oxidant with a reduction potential close to molecular oxygen (Kostka et al., 1995). Mn(III) is short-lived and unstable, but its stability is greatly increased when bound to ligands (Luther III et al., 2015). Ligand-bound Mn(III) is often the most abundant dissolved Mn species in sediment porewaters (Madison et al., 2013; Oldham et al., 2019) and soils (Heintze and Mann, 1947), with the potential to facilitate one-electron redox reactions in a variety of biogeochemical cycles (Luther III et al., 2015). Microbes accelerate the oxidation and reduction of Mn by orders of magnitude compared to abiotic mechanisms (Hem, 1963; Diem and Stumm, 1984; Morgan, 2005; Tebo et al., 2005; Learman et al., 2011; Luther et al., 2018; Jung et al., 2020). Yet, despite clear evidence for the environmental importance of Mn(III), knowledge about microbial Mn(III) cycling pathways remains fragmented.

To date, only *Shewanella* spp. (*Gammaproteobacteria*) have been confirmed to respire soluble Mn(III) (Kostka et al., 1995; Szeinbaum et al., 2014). *Shewanella* respire Mn(III) using the Mtr pathway (Szeinbaum et al., 2017), a porin-cytochrome (PCC) conduit that transports electrons across the periplasm for extracellular respiration of Mn(III/IV), Fe(III), and other metals (Richardson et al., 2012; Shi et al., 2016). Many Fe(II)-oxidizing *Betaproteobacteria* also contain PCCs (MtoAB, generally lacking the C subunit), which are proposed to oxidize Fe(II) to Fe(III) by running the PCC in reverse (Emerson et al., 2013; Kato et al., 2015; He et al., 2017). In some metal-reducing *Gammaproteobacteria* and *Deltaproteobacteria*, extracellular undecaheme (11-heme) UndA is thought to play a key functional role in soluble Fe(III) reduction (Fredrickson et al., 2008; Shi et al., 2011; Smith et al., 2013; Yang et al., 2013). UndA’s crystal structure shows a surface-exposed heme surrounded by positive charges, which may bind negatively-charged soluble iron chelates (Edwards et al., 2012). Environmental omics suggest that metal reduction by *Betaproteobacteria* may be widespread in the deep subsurface (Anantharaman et al., 2016; Hernsdorf et al., 2017). However, only a few Fe(III)-reducing *Betaproteobacteria* isolates have been characterized to date (Cummings et al., 1999; Finneran et al., 2003), and little is known about metal reduction pathways in *Betaproteobacteria*.

Manganese reduction coupled to methane (CH_4_) oxidation is a novel metabolism only recently discovered in cultures enriched in Archaea (Ettwig et al., 2016; Leu et al., 2020). Biological and geochemical evidence suggest that this metabolism may be found in a variety of environments (Beal et al., 2009; Crowe et al., 2011; Riedinger et al., 2014), including Fe-rich Lake Matano, Indonesia. In an attempt to explore whether CH4 can fuel microbial Mn(III) reduction in enrichments inoculated with sediments from Lake Matano, Indonesia, which has active and pronounced microbial Mn and CH_4_ cycles (Jones et al., 2011), we uncovered a novel betaproteobacterium as the most dominant and active member of our Mn(III)-reducing enrichment culture. Our results provide the first evidence for biological reduction of soluble Mn(III) outside *Gammaproteobacteria* and provide evidence for a new biochemical pathway involved in extracellular electron transfer.

## Results and discussion

### Enrichment of Mn(III)-reducing populations

Lake Matano, Indonesia, is a permanently stratified ultraoligotrophic lake (Crowe et al., 2008). Below its oxic surface waters, Lake Matano’s permanently anoxic and stratified waters are highly enriched in iron and manganese, and support the activity of Mn cycling organisms with organic carbon and CH_4_ as potential sources of electrons (Crowe et al., 2011; Jones et al., 2011; Kuntz et al., 2015; Sturm et al., 2019). We designed an enrichment strategy to select for microbes capable of anaerobic CH_4_ oxidation coupled to soluble Mn(III) reduction by incubating anoxic Lake Matano sediment communities with soluble Mn(III)-pyrophosphate as the electron acceptor (with 2% O_2_ in a subset of bottles), and CH4 as the sole electron donor and carbon source after pre-incubation to deplete endogenous organic carbon (see **Supporting Information** for enrichment details). Enrichment cultures were transferred into fresh media after Mn(III) was completely reduced to Mn(II), for a total of five transfers over 395 days. By the fourth transfer, cultures with CH_4_ headspace (with or without 2% O_2_) reduced ~80% of soluble Mn(III) compared to ~30% with N_2_ headspace (**Fig. 1**). 16S rRNA gene sequences were dominated by *Betaproteobacteria (Rhodocyclales;* 8-35%) and *Deltaproteobacteria (Desulfuromonadales;* 13-26%; **Fig. S1)**. ^13^CH_4_ oxidation to ^13^CO_2_ was undetectable (**Fig. S2**).

**Figure 1.**
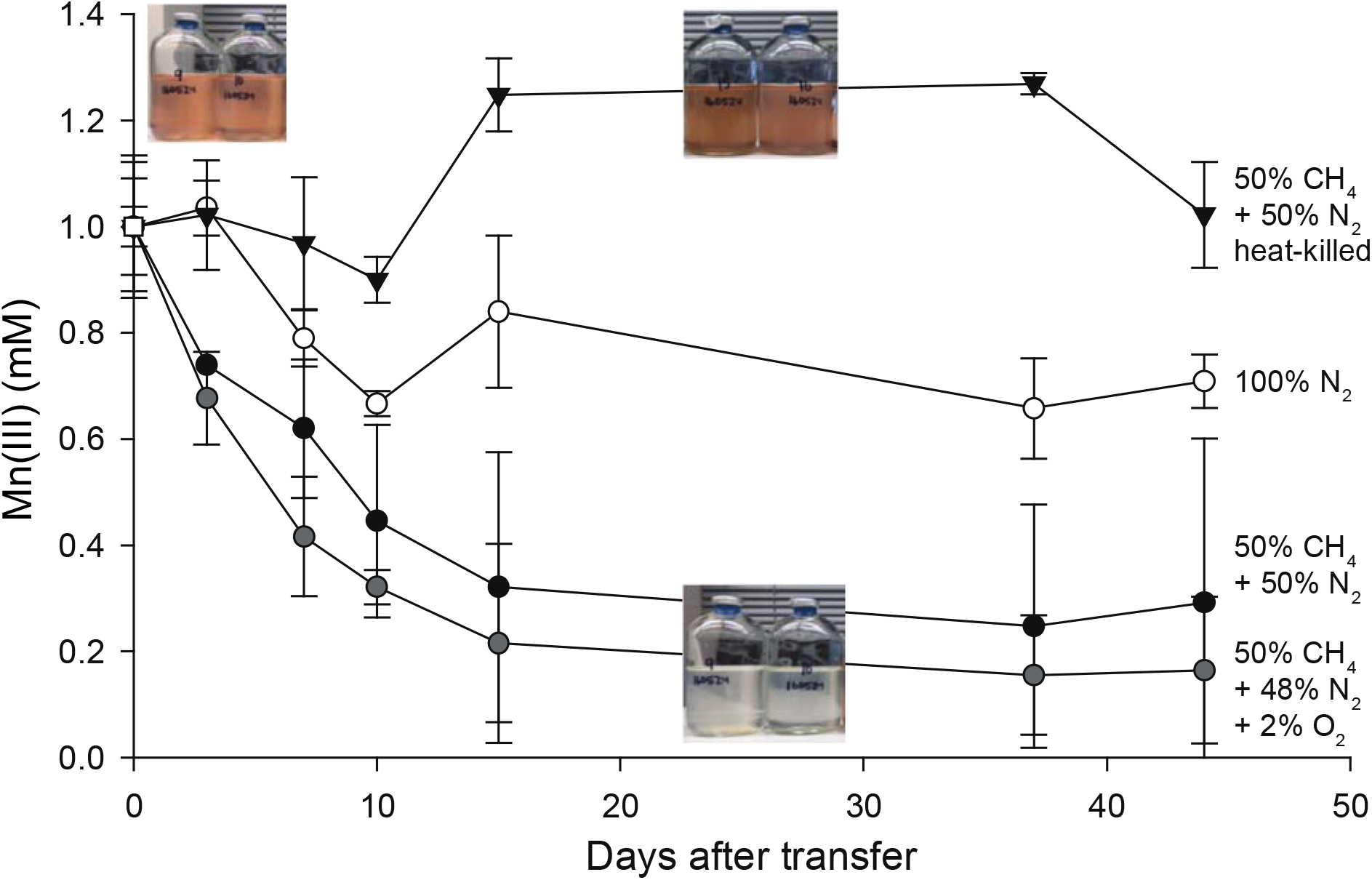
Consumption of Mn(III) in Lake Matano enrichments in the presence and absence of methane. Sediment-free cultures (transfer 4), from 335 days after the initial enrichment, were incubated for 45 days with 1 mM Mn(III) pyrophosphate as the sole electron acceptor. One set was incubated with Mn(III) and 2% O_2_. Initial bottle headspace contained 50% CH_4_ + 50% N_2_ (black circles), 50% CH_4_+48% N_2_+2% O_2_ (gray circles), 100% N_2_ (white circles), and 50% CH_4_+50% N_2_ heat killed controls (black triangles). Error bars are standard deviations from duplicate experiments. Color change from red to clear indicates Mn(III) reduction.

Samples for metagenomic and metaproteomic analysis were harvested from the fifth transfer (**Fig. 1; Fig. S1**). Out of 2,952 proteins identified in the proteome, 90% were assigned to *Betaproteobacteria;* of those, 72% mapped to a 99.5% complete metagenome-assembled genome (MAG; *Rhodocyclales* bacterium GT-UBC; NCBI accession QXPY01000000) with 81-82% average nucleotide identity (ANI) and phylogenetic affiliation to *Dechloromonas* spp. (**Table S1; Fig. S3**). This MAG is named here “*Candidatus* Dechloromonas occultata” sp. nov.; etymology: occultata; (L. fem. adj. ‘hidden’). The remaining 10% of proteins mapped to *Deltaproteobacteria;* of those, 70% mapped to a nearly complete MAG (*Desulfuromonadales* bacterium GT-UBC; NCBI accession RHLS01000000) with 80% ANI to *Geobacter sulfurreducens*. This MAG is named here “*Candidatus* Geobacter occultata”.

### Cytochrome expression during Mn(III) reduction

Cytochromes containing multiple *c*-type hemes are key for electron transport during microbial metal transformations, and therefore also expected to play a role in Mn(III) reduction. Numerous mono-, di-, and multi (>3)-heme cytochromes (MHCs) were expressed by “*Ca*. D. occultata” in Mn(III)-reducing cultures. Nine out of 15 MHCs encoded by the “*Ca*. D. occultata” MAG were expressed, including two decahemes similar to MtoA in Fe(II)-oxidizing *Betaproteobacteria* (**Tables 1, S2, S3; Figs. 2A, S4).** Several highly expressed MHCs were encoded on a previously unreported 19-gene cluster with 10 cytochrome-*c* proteins, hereafter *occA-S* (**Table 1; Figs. 2B, S5, S6**). OccP was predicted to be an extracellular undecaheme protein of ~100 kDa (922 amino acids). “*Ca*. Dechloromonas occultata” may reduce Mn(III) using the novel extracellular undecaheme OccP as the terminal Mn(III) reductase. Experimental verification of the function of the putative Occ complex is currently limited by the scarcity of genetically tractable *Betaproteobacteria*.

**Figure 2.**
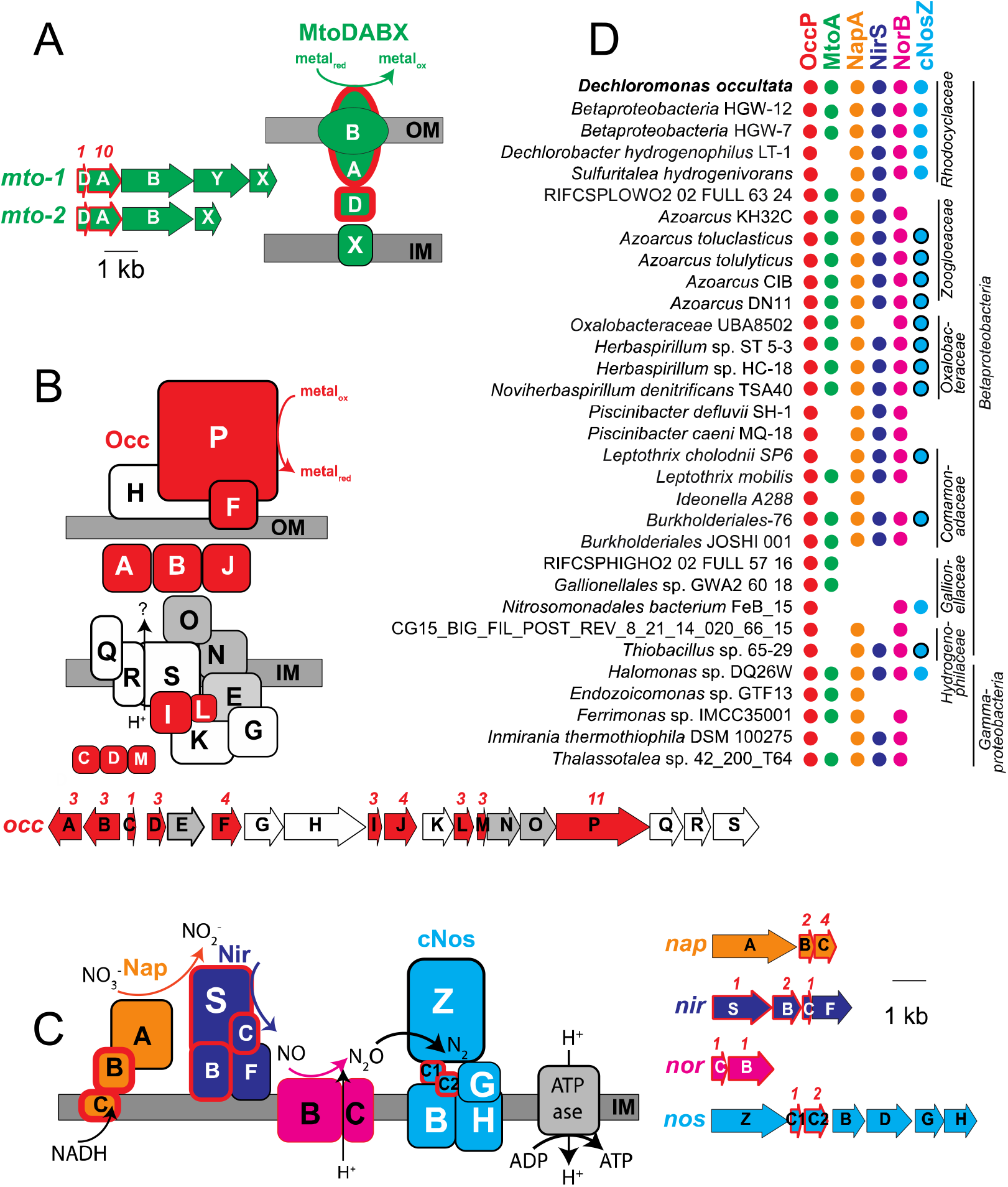
Gene arrangement, predicted protein location, and taxonomic distribution of major expressed respiratory complexes in “*Ca*. D. occultata”. **A:** MtoDAB(Y)X porin-cytochrome c electron conduit; **B:** OccA-S; **C:** denitrification complexes (Nap, Nir, Nor and cNos); **D:** Occurrence of key marker genes in *Betaproteobacteria* and *Gammaproteobacteria* with >95% complete genomes that encode OccP. Protein sequences from “*Ca*. D. occultata” were used as query against a genome database and searched using PSI BLAST. Matches with identities >40%, query coverage >80% and E values <10^-5^ were considered positive. Red fill around genes and proteins indicate cytochrome-*c* proteins. Black outlines around blue circles in D indicate type I nitrous oxide reductase to distinguish from blue dots (type II/cytochrome-nitrous oxide reductase). Gray-shaded genes on the *occ* gene cluster indicate 6-NHL repeat proteins. Protein locations shown are based on P-sort predictions. Numbers above genes indicate number of CxxCH motifs predicted to bind cytochrome *c*. IM: inner membrane; OM: outer membrane. For more details, see **Table 1** and **Table S3.**

**Table 1.**
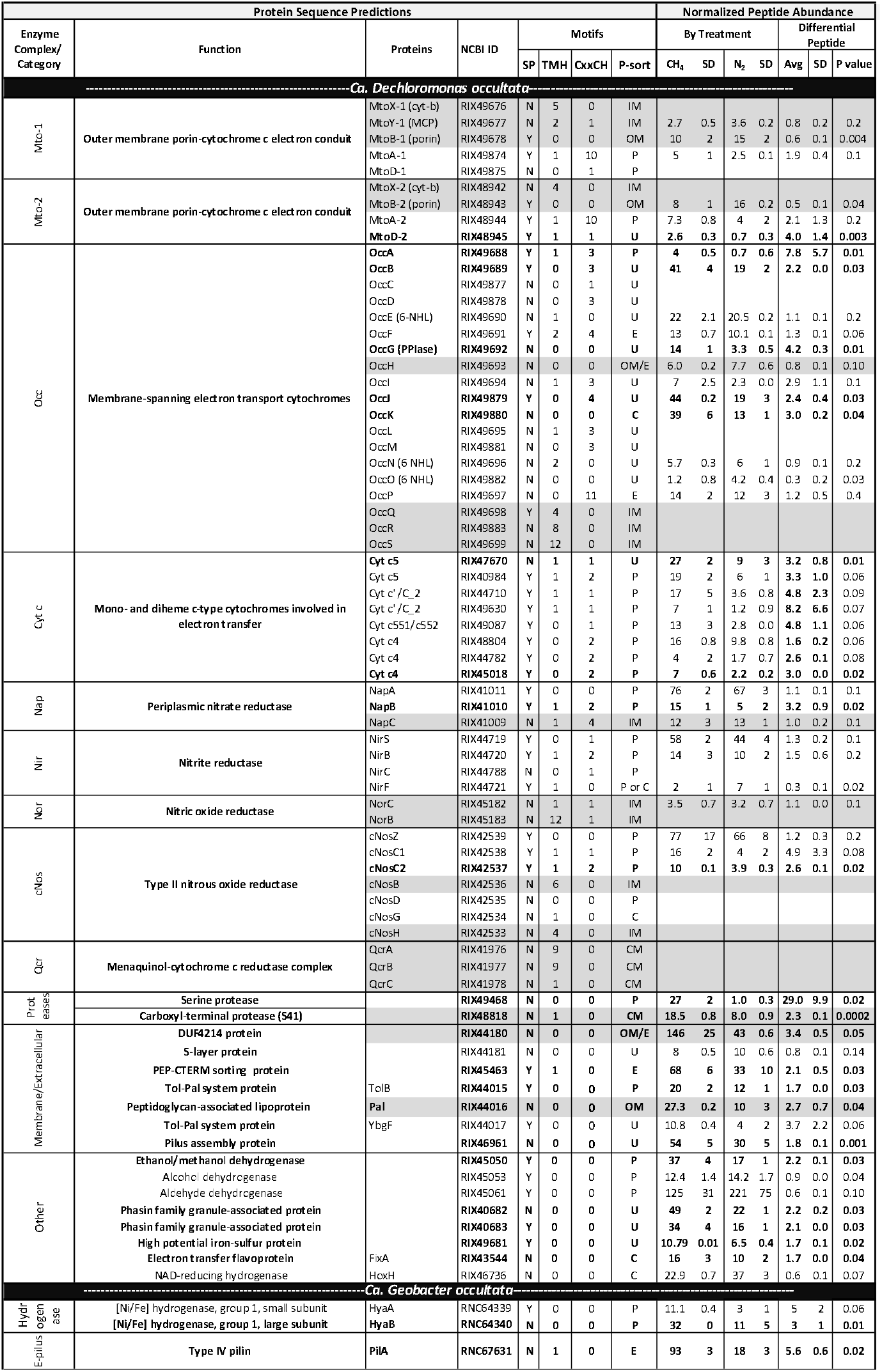
Expression levels for select “*Ca*. Dechloromonas occultata” and “*Ca*. Geobacter occultata” proteins in the presence of CH_4_ and N_2_. Gray boxes indicate membrane proteins^1^. Bold proteins indicate proteins that were significantly more expressed with CH_4_ than N_2_ (CH_4_/N_2_>1; p<0.05). P values indicate significance of abundance difference between CH_4_ and N_2_ treatments.

Proteins with 40-60% identity to the expressed “*Ca*. D. occultata” OccP protein were widely distributed in *Betaproteobacteria* from diverse freshwaters and deep subsurface groundwaters, as well as in several *Gammaproteobacteria* and one alphaproteobacterium (**Fig. 2D; Table S3).** Most *occP*-containing bacteria also possessed *mtoA* and denitrification genes (**Fig. 2D**; **Figs. S7**, **S8)**. These results widen the phylogenetic diversity of candidate extracellular MHCs that may be involved in microbial Mn(III) reduction.

### Heme-copper oxidases in “Ca. D. occultata”

“*Ca*. D. occultata” expressed high-affinity cbb_3_-type cytochrome c oxidase (CcoNOQP) associated with microaerobic respiration (**Table S4).** Features of the “*Ca*. D. occultata” *occS* gene product, including conserved histidine residues (H-94, H-411, and H-413) that bind hemes a and a_3_, as well as the H-276 residue that binds Cu_B_ (**Fig. S6**), suggest that OccS may function similarly to CcoN, the terminal heme-copper oxidase proton pump in aerobic respiration. All identified OccS amino acid sequences lack Cu_B_ ligands Y-280 and H-403, and most lack Cu_B_ ligands H-325 and H-326. OccS sequences also lack polar and ionizable amino acids that comprise the well-studied D and K channels involved in proton translocation in characterized cytochrome c oxidases (Blomberg and Siegbahn, 2014), but contain conserved H, C, E, D, and Y residues that may serve in alternate proton translocation pathways, similar to those recently discovered in qNOR (Gonska et al., 2018). OccS homologs were also found in *Azoarcus* spp. and deep subsurface *Betaproteobacteria* (**Fig. S6**).

### Expression of denitrification proteins and possible sources of oxidized nitrogen species

Periplasmic nitrate reductase (NapA), cytochrome nitrite reductase (NirS), and type II atypical nitrous oxide reductase (cNosZ; **Fig. S7**) were highly expressed by “*Ca*. D. occultata” **(Table 1)**. Expression of the denitrification pathway was not expected because oxidized nitrogen species were not added to the medium, to which the only nitrogen supplied was 0.2 mM NH_4_Cl (along with headspace N_2_). Oxidized nitrogen species could result from the oxidation of NH_4_Cl, but we did not find any of the canonical genes for aerobic nor anaerobic ammonia oxidation, nor did we measure any ammonium oxidation in experimental bottles from the transfer used to make **Figure 1**.

The expression of denitrification genes is controlled by a diverse array of transcriptional regulators that depend on different signals including low levels of oxygen, even in the absence of nitrate (Spiro, 2012; Lin et al., 2018). The close redox potential of Mn^3+^-pyrophosphate (~0.8 V; Yamaguchi and Sawyer, 1985)) to oxidized nitrogen species (0.35-0.75 V) at circumneutral pH and the lack of oxygen in the media could have induced the expression of denitrification genes simultaneously with Mn(III)-reduction genes. *Gammaproteobacteria*, for example, reduce Mn(III) even in the presence of nitrate (Kostka et al., 1995), and there is precedent for microbial use of multiple electron acceptors, e.g. “co-respiration” of oxygen and nitrate during aerobic denitrification (Chen and Strous, 2013; Ji et al., 2015).

Because solid-phase Mn(III) is known to chemically oxidize NH_4_^+^ (Aigle et al., 2017; Boumaiza et al., 2018), we tested for abiotic NH_4_^+^ oxidation by soluble Mn(III) (1 mM). Ammonium concentrations remained unchanged, and no N_2_O or NO_x_^-^ production was observed (**Fig. S8**), likely because our experiments lacked solid surfaces to mediate electron transfer. Similarly, N_2_O levels in the headspace of our experimental bottles with Mn(III)-reducing cultures were near or below the detection limit (data not shown). These findings are consistent with lack of detectable ammonium oxidation by Mn(III) pyrophosphate in estuarine sediments (Crowe et al., 2012).

### Electron donors

Methane was the only electron donor added intentionally to the enrichment cultures, to select for organisms that oxidize methane anaerobically. Yet, we did not detect ^13^CO_2_ after addition of ^13^CH_4_ (**Fig. S2**). One explanation is that ^13^CO_2_ was produced, but was subsequently assimilated by other members of the microbial community such as abundant Deltaproteobacteria **(Fig. S1)**, as observed in previous studies (Wegener et al., 2008). A filtration step included in our protocol to measure ^13^CO_2_ would have excluded ^13^C-enriched biomass from our analyses. Alternatively, we considered other electron donors that might have been unintentionally present in trace amounts, but sufficiently abundant to drive the observed ~300-600 μM Mn(III) reduction **(Fig. 1)**. The ethanol catabolism pathway (PQQ-dependent methanol/ethanol dehydrogenase (RIX45050), quinoprotein alcohol dehydrogenase (RIX45053), and an NAD^+^-dependent aldehyde dehydrogenase-II (RIX45061)) were all highly expressed in “Ca. *D. occultata*” **(Table 1)**. Ethanol could have been introduced to the bottles during culture preparation during sterilization of bottle stoppers. Based on the stoichiometry of ethanol oxidation coupled to Mn(III) reduction:

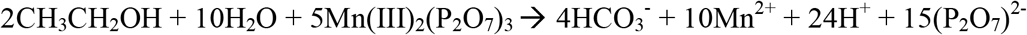

150 μM ethanol would be required to reduce 600 μM of Mn(III), which equates to ~1 μL of 70% ethanol (12 M) into 100 mL culture medium. We conclude that trace contamination of ethanol was likely the major electron donor to our cultures.

It is also possible that other substrates, such as H_2_ from fermentation by other microbes in the enrichment or from impurities in the headspace gas, could have supplied another source of electrons. Indeed, an NAD-reducing hydrogenase (RIX44099-100) was expressed by “*Ca*. D. occultata” (**Table 1**). Based on the stoichiometry of H_2_ oxidation coupled to Mn(III) reduction:

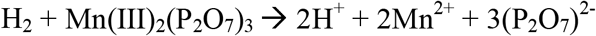

600 μM H_2_ would be required to reduce 600 μM of Mn(III). Thus, H_2_ may have contributed electrons to Mn(III) reduction, but is not likely sole electron donor. A combination of ethanol, H_2_, and other trace contaminants would likely have been necessary to provide enough electrons for the additional reduction of Mn(III) observed in the ^13^CH_4_-amended cultures compared to the controls lacking ^13^CH_4_. There is precedent for other metal-reducers simultaneously using H_2_ and an organic electron donor (Brown et al., 2005).

Another trace source of organics to our cultures could have been leaching from the rubber stoppers, which were black bromobutyl and pre-boiled in 0.1 N NaOH. A previous study reported that organics leaked an array of n-alkanes (C_16_–C_34_) and unidentified organic contaminants in black bromobutyl stoppers (Niemann et al., 2015). It is also conceivable that trace organic was introduced as impurities in solid Mn(III) oxide powder (99% purity) used to synthesize Mn(III)-pyrophosphate.

Finally, we considered the possibility that 0.2 mM NH_4_^+^, added to the cultures as a nitrogen source, could have provided the electron donor, via an unknown pathway. Based on the stoichiometry of NH_4_^+^ oxidation coupled to Mn(III) reduction:

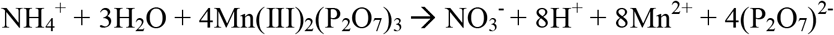

0.2 mM NH_4_^+^ would supply 1.6 mM electron equivalents, which is more than enough to account for the observed reduction of 600 μM of Mn(III). This process could operate cryptically if the oxidized products were reduced to N_2_ via denitrification enzymes, such as nitrous oxide reductase (cNosZ), which was one of the most abundant proteins expressed in Mn(III)-reducing cultures **(Fig. 2c, Table 1).**

### Carbon metabolism

“*Ca*. D. occultata” appeared to be growing mixotrophically. “Ca. *D. occultata”* encoded several central metabolic pathways, including a complete TCA cycle with a glyoxylate bypass, an incomplete (acetate-dependent) 3-hydroxypropionate bicycle (3-HP), a modified Calvin-Benson-Bassham (CBB) pathway, and a pathway for synthesis of polyhydroxybutyrate **(Fig. S11)**. In addition, “Ca. *D. occultata”* encoded genes for organic carbon transport, and lactate, acetate, and propionate utilization **(Fig. S11)**. Like *D. agitata* and *D. denitrificans*, the CBB pathway of “*Ca*. D. occultata” did not encode RuBisCO and sedoheptulose-1,7-bisphosphatase (SHbisPase; **Fig. S10)**; SHbisPase may be replaced by 6-phosphofructokinase and an energy-generating pyrophosphatase (RIX41248; Kleiner et al., 2012; Zorz et al., 2018). The presence of incomplete carbon fixation pathways and for organic carbon utilization pathways suggests that “Ca. *D. occultata”* relies on organic carbon to fix inorganic carbon mixotrophically. The source of this organic carbon could have been ethanol, which is converted to acetate via the pathway discussed in the previous section.

### Effect of methane

Although we did not measure appreciable ^13^CH_4_ oxidation to ^13^CO_2_, CH_4_ stimulated Mn(III) reduction and cytochrome expression in “*Ca*. D. occultata” enrichment cultures. While the specific role of CH_4_ in Mn(III) reduction remains unknown, the addition of CH_4_ appeared to significantly stimulate expression of many cytochrome *c* proteins, including OccABGJK, MtoD-2, and cytochrome-*c*4 and -*c*5 proteins associated with anaerobic respiration (p < 0.05; **Table 1; Fig. 2C**). Expression of several “*Ca*. D. occultata” proteins involved in outer membrane structure and composition - including an extracellular DUF4214 protein located next to an S-layer protein similar to those involved in manganese binding and deposition (Wang et al., 2009), a serine protease possibly involved in Fe(III) particle attachment (Burns et al., 2009), an extracellular PEP-CTERM sorting protein for protein export (Haft et al., 2006), and a Tol-Pal system for outer membrane integrity - was higher in the presence of CH_4_ (**Table 1)**.

### Transporters and sensors

Numerous transporters were present in the “*Ca*. D. occultata” genome, including 26 TonB-dependent siderophore transporters, 13 TRAP transporters for dicarboxylate transport, as well as ABC transporters for branched-chained amino acids and dipeptides and polypeptides (**Table S4**). “*Ca*. D. occultata” also contained a large number of environmental sensing genes: 52 bacterial hemoglobins with PAS-PAC sensors, 8 TonB-dependent receptors, and 8 NO responsive regulators (Dnr: Crp/fr family; **Table S4**). Uniquely in “*Ca*. D. occultata”, PAC-PAS sensors flanked accessory genes *nosFLY* on the *c-nosZ* operon (**Fig. S7**). Comparison of these flanking PAC-PAS sensors in “*Ca*. D. occultata” with O_2_-binding sensors revealed that an arginine ~20 aa upstream from the conserved histidine as the distal pocket ligand for O_2_-binding is not present in either sensor (**Fig. S11**), suggesting that the sensor may bind a different ligand, possibly NO, consistent with the placement of these genes next to cNosZ (Shimizu et al., 2015).

### Nutrient storage

Active synthesis of storage polymers suggested that “*Ca*. D. occultata” was experiencing electron acceptor starvation at the time of harvesting, consistent with Mn(III) depletion in the bottles (Liu et al., 2015; Guanghuan et al., 2018). Polyphosphate-related proteins, including phosphate transporters, polyphosphate kinase, polyphosphatase, and poly-3-hydroxybutyrate synthesis machinery were detected in the proteome (**Table S4**). Polyphosphate-accumulating organisms store polyphosphates with energy generated from organic carbon oxidation during aerobic respiration or denitrification. These stored compounds are later hydrolyzed when respiratory electron acceptors for ATP production are limiting. Cyanophycin was actively synthesized for nitrogen storage.

### Geobacter

“*Ca*. Geobacter occultata” expressed proteins in the TCA cycle at moderate abundance. “*Ca*. G. occultata” contained 17 multiheme c-type cytochromes, none of which were detected in the proteome. The lack of expression of electron transport and metal-reducing pathways makes it unlikely that “*Ca*. G. occultata” was solely responsible for Mn(III) reduction observed in the incubations. A periplasmic group I Ni-Fe hydrogenase (RNC64340; 91% identity to a protein (RLB64899) from *Geobacter* MAG from terrestrial hot spring sediment) and a type IV pilin (RNC67631; 10% aromatics, 87% identity to *Geobacter pickeringii* (Holmes et al., 2016)) were significantly more expressed in the presence of CH_4_ than N_2_ in the “*Ca*. G. occultata” proteome (p < 0.05; **Table 1**). It is possible that “*Ca*. G. occultata” transferred electrons to “*Ca*. D. occultata” via e-pilins (e.g. direct interspecies electron transfer), contributing to the higher rates of Mn(III) reduction in the presence of CH_4_ vs. N_2_. The possible involvement of *Geobacter* e-pilins in Mn(III) reduction remains an open question, due to the lack of studies examining the possibility of Mn(III) reduction in *Deltaproteobacteria*.

### Conclusions

To our knowledge, this study provides the first evidence for biological reduction of soluble Mn(III) by a bacterium outside of the *Gammaproteobacteria* class. The dominant bacterium in Mn(III)-reducing enrichment cultures was “*Ca*. D. occultata”, a member of the *Rhodocyclales* order of *Betaproteobacteria*. “*Ca*. D. occultata” expressed decahemes similar to the Mto pathway, and *occ* genes, including a novel extracellular undecaheme (OccP), which are predicted to encode a new respiratory electron transport pathway. The novel *occ* operon was found to be widespread in *Betaproteobacteria* from the deep subsurface, where metal cycling can fuel microbial metabolism. We also found highly expressed peptides from various central metabolic cycles and organic substrate utilization pathways, suggesting that “Ca. *D. occultata”* may have been using multiple pathways simultaneously for energy generation and carbon assimilation during Mn(III) reduction.

Puzzles remain about whether “*Ca*. D. occultata” can transform two potent greenhouse gases: methane and nitrous oxide. Although “*Ca*. D. occultata” was enriched with CH_4_ as the sole electron donor and cultures reduced Mn(III) more rapidly in the presence of CH_4_, no CH_4_ oxidation activity was measured in Mn(III)-reducing cultures, and proteomic data suggested that “*Ca*. D. occultata” was growing mixotrophically rather than assimilating CH_4_. Further, although we did not add oxidized nitrogen compounds to our media, and Mn(III) did not chemically oxidize NH_4_^+^ under our culture conditions, type II nitrous oxide reductase (cNosZ) was one of the most abundant proteins expressed in Mn(III)-reducing cultures. The role of cNosZ and other denitrification enzymes in “*Ca*. D. occultata” metabolism, and their possible connection to Mn(III) reduction, remain to be investigated.

## Acknowledgements

This research was funded by NASA Exobiology grant NNX14AJ87G. Support was also provided by a Center for Dark Energy Biosphere Investigations (NSF-CDEBI OCE-0939564) small research grant and supported by the NASA Astrobiology Institute (NNA15BB03A) and a NASA Astrobiology Postdoctoral Fellowship to NS. SAC was supported through NSERC CRC, CFI, and Discovery grants. We thank Marcus Bray, Andrew Burns, Caleb Easterly, Ellery Ingall, Pratik Jagtap, Cory Padilla, Angela Peña, Johnny Striepen, Yael Toporek, and Rowan Wolschleger for technical assistance. We thank Karen Lloyd, Nagissa Mahmoudi, and Emily Weinert for helpful discussions.

## Footnotes

1 SP: signal peptide (Y:present/N:absent); TMH: numbers of transmembrane helices; CxxCH: number of heme-binding motifs; P-sort: predicted cellular location based on Psortb v.3.0. MCP: methyl-accepting chemotaxis protein; PPIase: Peptidyl-proline isomerase; P: periplasm, C: cytoplasm; OM: outer membrane; IM: inner membrane, E: extracellular; U: unknown. MtoX and MtoY were predicted to be an inner membrane cytochrome-b protein and a methyl-accepting chemotaxis protein, respectively. Membrane proteins may be underrepresented by mass spectrometry-based metaproteomic analyses, which inherently favor soluble over insoluble membrane-bound or hydrophobic proteins.

