## Supporting Information for "Novel insights into the taxonomic diversity and molecular mechanisms of bacterial Mn(III) reduction"

**Running title:** Novel undecaheme in Betaproteobacteria

**Content:**

1. **Experimental procedures**
2. **Supplemental tables**
3. **Supplemental figures**
1. **Experimental procedures**

**Source of inoculum.** Lake Matano is a metal-rich, ancient ocean analog with an active Mn cycle (1, 2). Organic carbon in Lake Matano is mostly mineralized via methanogenesis (3). A 15-cm sediment core from 200 m water depth in Lake Matano, Sulawesi Island, Indonesia (02°26′27.1′′S, 121°15′12.3′′E; *in situ* sediment temperature ~27°C) was sampled in November 2014 and sub-sampled at 2.5-cm increments. Sediments were sealed in gas-tight Mylar bags with no headspace (4) and stored at 4°C for ~1 year.

**Inoculation of enrichment cultures.** Mylar bags containing sediment samples were opened in an anoxic chamber (97% N_2_ and 3% H_2_; Coy Laboratory Products, Grass Lake, MI, USA). Sediments from each 2.5 cm subsample were transferred to 160 mL serum bottles, diluted 1:2 with minimal media, and pre-incubated for 45 days at 30°C in 100% N_2_ headspace to deplete endogenous organic carbon, electron donors, and electron acceptors. Sediments from the top 5 cm were subsequently mixed together and transferred to defined medium at a 1:20 dilution (transfer 1, day 45) amended with Mn(III) and a headspace of CH_4_:N_2_ (50:50) or N_2_. Subsequent transfers were carried out in the same way (transfer 2, day 91; transfer 3, day 183; transfer 4, day 230) for CH_4_ headspace cultures, with heat-killed and substrate controls generated each time using the newly transferred culture (10% v/v dilution). By day 210, enrichments appeared to be sediment-free, except for microparticles. The fifth transfer (day 245) inoculated using non-labeled methane, was used entirely for metaproteomic analysis after visual confirmation of active Mn(III) reduction in the live enrichment bottles (see **Figs. 1**, **S1**). Two 100 mL bottles were pooled together two obtain 200 mL duplicates for each treatment, centrifuged (10,000 x g, 30 min, 4°C) and supernatant-free pellets were stored at -80°C until protein extraction and metaproteomic sequencing**.**

Defined medium consisted of modified artificial freshwater medium lacking nitrate and sulfate, developed based on the pore water composition of Lake Matano sediments as described in prior work (5). The medium contained 3 mM NaHCO_3_, 825 μM MgCl_2_, 550 μM CaCO_3_, 225 μM NH_4_Cl, 5 μM Na_2_HPO_4_, 3.5 μM K_2_HPO_4_, and a trace metal solution (1 nM CuCl_2_, 1.5 nM Na_2_MoO_4_, 2.5 nM CoCl_2_, 23 nM MnCl_2_, 9 nM FeCl_3_, 4 nM ZnCl_2_, 0.091 μg/L vitamin B_12_, 0.091 μg/L biotin, and 18.18 μg/L thiamine. Vitamins were filter-sterilized and added after autoclaving. Bottles were stoppered with sterile black bromobutyl stoppers (Geo-Microbial Technologies, Ochelata, OK, USA; pre-boiled in 0.1 N NaOH), and crimped with aluminum seals. An acetate-free 10 mM stock of Mn(III)-pyrophosphate was prepared using solid Mn_2_O_3_ (99% purity, 325 mesh powder, Sigma Aldrich) instead of Mn(III)-acetate (6) and filter-sterilized. Mn(III)-pyrophosphate was added at a final concentration of 1 mM. Bottles were purged with 99.9% N_2_ for 20 min, and were appropriate, CH_4_ was injected with a 50% headspace volume of CH_4_ at a 1:1 labeled to unlabeled ratio (99.9% CH_4_ and 99% ^13^CH_4_; Cambridge Isotope Laboratories, Tewksbury, MA, USA). Heat-killed controls were autoclaved prior to Mn(III) or CH_4_ addition. All treatments were duplicated, and bottles were incubated in the dark at 30°C.

**Substrate utilization.** The benzidine method was used to measure Mn(III) consumption (7) throughout the transfer 4 enrichment. Methane (^13^CH_4_) oxidation was monitored by measuring ^13^C enrichment in dissolved inorganic carbon as described in (5).

**16S rRNA gene amplicon sequencing.** To identify the dominant microbial community members, we analyzed the microbial community composition of samples taken at the end of each enrichment period by sequencing 16S rRNA gene amplicons as described previously (5). Reads were analyzed using Mothur (8) following its MiSeq standard operating procedure (<https://www.mothur.org/wiki/MiSeq_SOP>, accessed November 2017). Merged reads were dereplicated and aligned to the ARB SILVA SSU database release 123 (July 23, 2015). Homopolymers longer than 8bp were filtered out. Reads were then clustered into OTUs at 97% similarity based on uncorrected pairwise distance matrices. OTUs were classified using the ARB SILVA SSU reference taxonomy database release 123.

**Metagenome (DNA) sequencing and assembly.** Community DNA was processed using the Nextera XT DNA Sample Prep kit and sequenced using a paired-end Illumina MiSeq 600 kit. Raw reads were submitted to NCBI. The accessions for the study and samples in the submission are PRJNA489678, LakeMatanoMn3_Enrichment (SAMN10343573). The accession numbers for the N_2_ headspace experiment and run are LM_Mn(III)_2018 (SRX5007804) and LakeMatano_11_NoMethane_R1.fastq.gz (SRR8188020), and the accession numbers for the CH_4_ headspace experiment and run are LM_Mn(III)_CH4_2018 (SRX5007805) and LakeMatano_9_Methane_R1.fastq.gz (SRR8188019).

Barcoded sequences were de-multiplexed, trimmed (length cutoff 100 bp), and filtered to remove low quality reads (average Phred score <25) using Trim Galore! (<http://www.bioinformatics.babraham.ac.uk/projects/trim_galore/>). Forward and reverse reads were assembled using SPAdes (9) with the ‘meta’ option. Metagenomic reads were deposited in NBCI. Contigs ≥ 500 nt were organized into MAGs based on tetranucleotide frequency and sequence coverage using MaxBin 2.0 (10). MAG completeness and contamination were estimated by lineage-specific marker genes using CheckM (11). We obtained one *Betaproteobacteria* metagenome-assembled genome (MAG; *Rhodocyclales* bacterium GT-UBC, NCBI accession QXPY01000000) with 99.53% completeness, 0.02% contamination, 60.9% GC content, and 4,555 protein-coding genes. We provisionally classified *Rhodocyclales* bacterium GT-UBC as a new species within the *Dechloromonas* genus, which we named “*Candidatus* Dechloromonas occultata” sp. nov.; etymology: occultata; (L. fem. adj. ‘hidden’), based on its resistance to cultivation. We also obtained one *Deltaproteobacteria* MAG (*Desulfuromonadales* bacterium GT-UBC; NCBI accession RHLS01000000) with 99.36% completeness, 0.64% contamination, 59.9% GC content, 3,617 protein-coding genes*,* and 80% ANI to *Geobacter sulfurreducens.* We provisionally named *Desulfuromonadales* bacterium GT-UBC “*Candidatus* Geobacter occultata” sp. nov.

**Comparative genomic analysis.** The “*Ca*. D. occultata” MAG was annotated using RAST. Metabolic pathways were identified in RAST (12-14), and manually checked for completeness in PATRIC (15), following pathways reported in KEGG (https://www.genome.jp/kegg/) and MetaCyc (https://metacyc.org/) and confirming potential new variants in the literature (references where appropriate throughout the manuscript). For incomplete pathways, the three other genomes of isolated *Dechloromonas* strains (*D. aromatica*, *D. denitrificans*, and *D. agitata*) in PATRIC were searched for missing proteins. Central metabolic and secondary pathways were compared among *Dechloromonas* spp. and other metal-cycling species using the RAST subsystems option, searching for specific annotated proteins with RAST, and within PATRIC using the Compare Region Viewer and its heatmap options. Proteins of interest were characterized *in silico* based on conserved domains and homology searches with BLAST and NCBI tools. Localization of proteins was predicted using PSORTb model ECSVM (16). The synteny of gene clusters containing functional genes of interest were analyzed using Simple Synteny (<https://www.dveltri.com/simplesynteny/cfinder.html>, last accessed July 2018). Putative multiheme *c*-type cytochromes (≥3 Cxx(x)CH motifs) were identified using a previously reported Python script (<https://github.com/bondlab/scripts>, (17)).

**Phylogenetic analysis.** The evolutionary history of functional genes was inferred using MEGA7 (18) with the Maximum Likelihood method based on the JTT matrix-based model (19). After all gaps were eliminated, initial tree(s) for the heuristic search were obtained automatically by applying Neighbor-Join and BioNJ algorithms to a matrix of pairwise distances estimated using a JTT model, and then selecting the topology with superior log likelihood value. The trees were drawn to scale, with branch lengths measured in the number of substitutions per site and bootstrap values based on 500 replicates. The synteny of selected genes was determined using Simple Synteny (20). The evolutionary history of selected whole genomes of Proteobacteria was reconstructed using GToTree (21) with genomic NCBI IDs as input, retrieved manually. Single-copy genes (SCGs) were identified from a set of 74 single-copy genes. GToTree uses concatenated alignments of identified SCGs to build the phylogenomic tree with FastTree. The final tree was viewed and edited in FigTree V1.4.4 (22).

**Protein digestion and desalting.** To solubilize oxidized metal precipitants and precipitate proteins, 100 µL of 20% trichloroacetic acid (4°C) was added to each sample and incubated on ice for 1 hour. Bacterial cells and soluble proteins were pelleted at 10,000 x g (1 hr, 4°C). Cells were then resuspended in 100 µL of 6 M urea in 50 mM NH_4_HCO_3_ and lysed using a sonicating probe (3 watts; 15s, 5 times), alternating in dry ice in ethanol to keep the sample cold. Sonication, digestion, and desalting proceeded as previously described (23). Briefly, after sonication and protein quantification using the Bradford assay (Bio-Rad, Hercules, CA), tris(2-carboxyethyl)phosphine (TCEP) was added to reduce samples (1 hr, 37°C), and iodoacetamide was used as the alkylating agent (1hr, in dark, RT). NH_4_HCO_3_ and HPLC-grade methanol were added to each sample to dilute the urea to allow the trypsin digestion to proceed. Trypsin was added in a 1:20 ratio and incubated overnight at RT. The digestion was stopped by adding small aliquots of 10% formic acid until a pH < 2 was achieved. Prior to desalting the peptides, samples were dried down and reconstituted in 5% acetonitrile with 0.1% trifluoroacetic acid. Desalting was carried out with MicroSpin C18 columns following the manufacturer’s instructions (The Nest Group). Peptides were dried and reconstituted in 5% ACN with 0.1% formic acid to achieve concentrations of 2 µg µL^-1^.

**LC-MS/MS.** The mass spectrometry analysis was performed on a QExactive at the University of Washington Proteomics Resource (Seattle, WA). Samples were separated and introduced into the mass spectrometer (MS) by reverse-phase chromatography using a Manufactured PicoTip fused silica capillary column (30 cm long, 75 µm i.d.) packed with C18 particles (Dr. Maisch ReproSil-Pur; C18-Aq, 120 Å, 3 µm) fitted with a 3 cm long, 100 µm i.d. precolumn (Dr. Maisch ReproSil-Pur; C18-Aq, 120 Å, 3 µm). Peptides were eluted using an acidified (formic acid, 0.1% v/v) water-acetonitrile gradient (5–35% acetonitrile in 90 min) and mass spectrometry was performed on a Thermo Fisher (San Jose, CA) QExactive (QE). The top 20 most intense ions were selected for MS2 acquisition from precursor ion scans of 400–1200 m z^−1^. Centroid full MS resolution data was collected at 70,000 with AGC target of 1E6 and centroid MS2 data was collected at resolution of 35,000 with AGC target of 5E4. Dynamic exclusion was set to 15 seconds and +2, +3, +4 ions were selected for MS2 using data dependent acquisition mode (DDA). Quality control (QC) peptide mixtures were analyzed every fifth injection to monitor chromatography and MS sensitivity. Skyline was used to determine that QC standards did not deviate >10% through all analyses (24). The mass spectrometry data are deposited to the ProteomeXchange Consortium via the PRIDE partner repository with the dataset identifier PXD011642 (username:; password: PFsKqEHP).

**Protein identification and data analyses.** Peptide identifications from mass spectrometry data were completed using Comet (25). The protein database used for correlating spectra with protein identifications was generated from the metagenome by Prokka (26), and from each individual bin using RAST (13) and included the MAGs to improve peptide spectra correlations (15). This was then combined with 50 common contaminants and the QC peptides. Comet parameters included: reverse concatenated sequence database search, trypsin enzyme specificity, cysteine modification of 57 Da (resulting from the iodoacetamide) and modifications on methionine of 15.999 Da (oxidation). Concatenated target–decoy database searches were completed and minimum protein and peptide thresholds were set at P > 0.95 on ProteinProphet and P > 0.99 on PeptideProphet (27). Protein identifications from the whole-cell lysates were accepted by ProteinProphet if the above mentioned thresholds were passed, two or more peptides were identified (PeptideProphet), and at least one terminus was tryptic (27). Calculated false discovery rates (FDR) were <0.01. Resulting data files were combined and normalized spectral abundances were calculated in QPROT with Abacus (28). Abacus parameters include initial probability threshold of 0.5 on peptides, and a minimum protein probability of 0.8. Abacus provides consistent protein inferences across biological and technical replicates. Abacus spectral abundance outputs were analyzed with QSpec, a statistical framework within QPROT, to determine log fold changes between treatments. Log fold change in protein abundances calculated using QSpec were accepted if ≥0.5 and Zstatistic score ≥2.0 (increased abundance across all replicates) or <-0.5 and Zstatistic score <-2.0 (decreased abundance across all replicates) (28). Peptide counts were normalized to total peptide counts for each treatment. Averages of normalized technical replicates were used to compare treatments with and without methane. A two-tailed paired t-test was carried out using Excel to test the null hypothesis of no differential expression among treatments and determine the p-value associated with each change.

**Cultivation attempts.** Isolation strategies were designed considering the metabolic potential of “*Ca.* D. occultata” but failed to isolate the targeted organism. Samples from highly enriched cultures were inoculated with acetate for denitrification or microaerobic Fe(II) oxidation. For Fe(II) oxidation, samples were inoculated into two-layered FeS vs. O_2_ gradient tubes (29), with 1 mM acetate in the top layer. There was no Fe(II) oxidation in Mn(III)-amended treatments. With O_2_ addition, visual evidence for Fe(II) oxidation was observed, and a *Comamonas* spp. with closest hits to environmental sequences from lake sediment was isolated. With acetate and Mn(III), a [*Comamonas aquatica*](https://blast.ncbi.nlm.nih.gov/Blast.cgi#alnHdr_1049746289) strain was isolated.

1. **Supplemental Tables**

**Table S1. Average nucleotide identity of *Dechloromonas* species.** Numbers in the table indicate percentage of whole genome nucleotide identity.

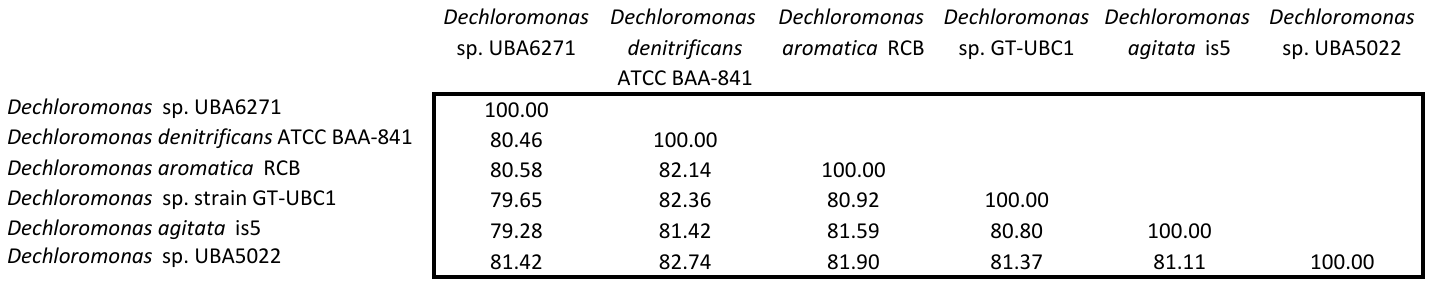

**Table S2. List of multiheme cytochromes encoded by “*Candidatus* Dechloromonas occultata”.** Those in bold indicate expressed proteins (see **Table 1**).

| **NCBI ID** | **Predicted function** | **Heme-binding motifs** | **Protein length (aa)** |
| --- | --- | --- | --- |
| **RIX41009** | **NapC-like** | **4** | **207** |
| RIX43626 | NapC-like | 4 | 198 |
| **RIX48944** | **MtoA-2** | **10** | **320** |
| **RIX49874** | **MtoA-1** | **10** | **331** |
| RIX49876 | Hypothetical protein | 3 | 127 |
| **RIX49688** | **OccA** | **3** | **327** |
| **RIX49689** | **OccB** | **3** | **343** |
| RIX49878 | OccD | 3 | 175 |
| **RIX49691** | **OccF** | **4** | **289** |
| **RIX49694** | **OccI** | **3** | **139** |
| **RIX49879** | **OccJ** | **4** | **338** |
| RIX49695 | OccL | 3 | 194 |
| RIX49881 | OccM | 3 | 87 |
| **RIX49697** | **OccP** | **11** | **922** |
| RIX49727 | Hypothetical protein | 5 | 695 |

**Table S3. Genomes containing MtoA, OccP, NapA, NirS, NorB, or cNosZ homologs in Alpha-, Beta-, and Gammaproteobacteria**. “Ecosystem Type” refers to the source of inoculum for pure cultures or the source of environmental DNA for assembled genome of uncultured organisms. Spreadsheet is attached as supplemental file.

**Table S4. Expression levels for “*Ca*. D. occultata” proteins during Mn(III) reduction with and without CH_4_.** Peptide counts are normalized to total “*Ca*. D. occultata” proteins x 10,000. Blank cells indicate proteins with <5 normalized peptide counts. Gray boxes indicate membrane proteins with that may be underrepresented by proteomic analyses. SP: signal peptide (Y:present/N:absent); TMH: numbers of transmembrane helices; # CxxCH: number of heme-binding motifs; P-sort: predicted cellular location. P: periplasm, C: cytoplasm; OM: outer membrane; IM: inner membrane, E: extracellular; U: unknown. Spreadsheet is attached as supplemental figure.

2. **Supplemental Figures**

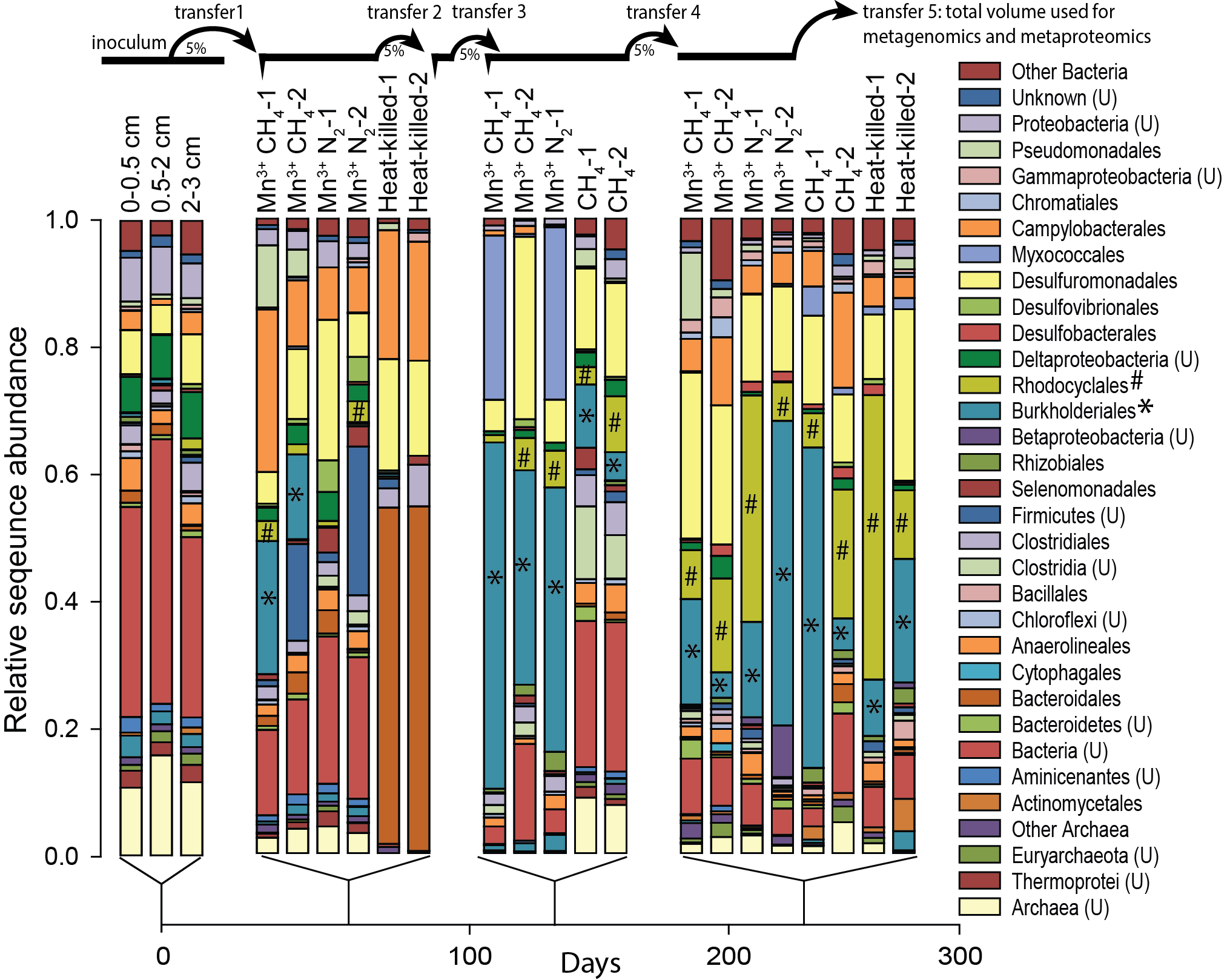

**Figure S1.** **Taxonomic succession in enrichment culture.** Relative abundance of taxa enriched from samples from Lake Matano sediments over a 335-day period, based on ~200 bp 16S rRNA gene amplicon sequences. Only live treatments with CH_4_ and Mn(III) were transferred. (U) indicates unclassified taxa.

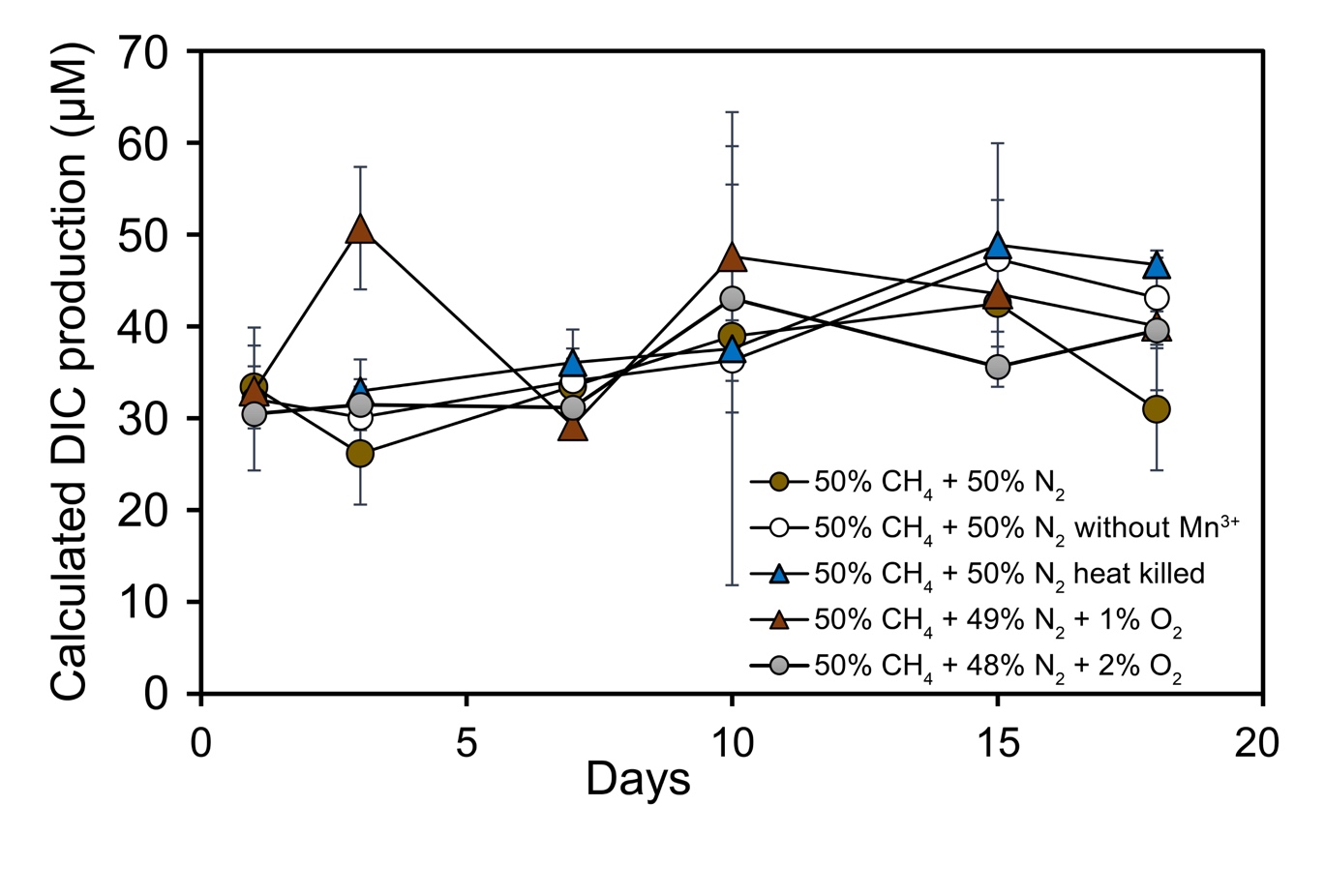

**Figure S2. Methane oxidation after the fourth transfer of enrichment cultures.** This graph shows the concentration of methane-derived dissolved inorganic carbon (DIC) in sediment enrichments amended with ^13^CH_4_, calculated based on isotopic enrichment values and total DIC based on (30). Errors bars represent standard deviation of duplicate measurements.

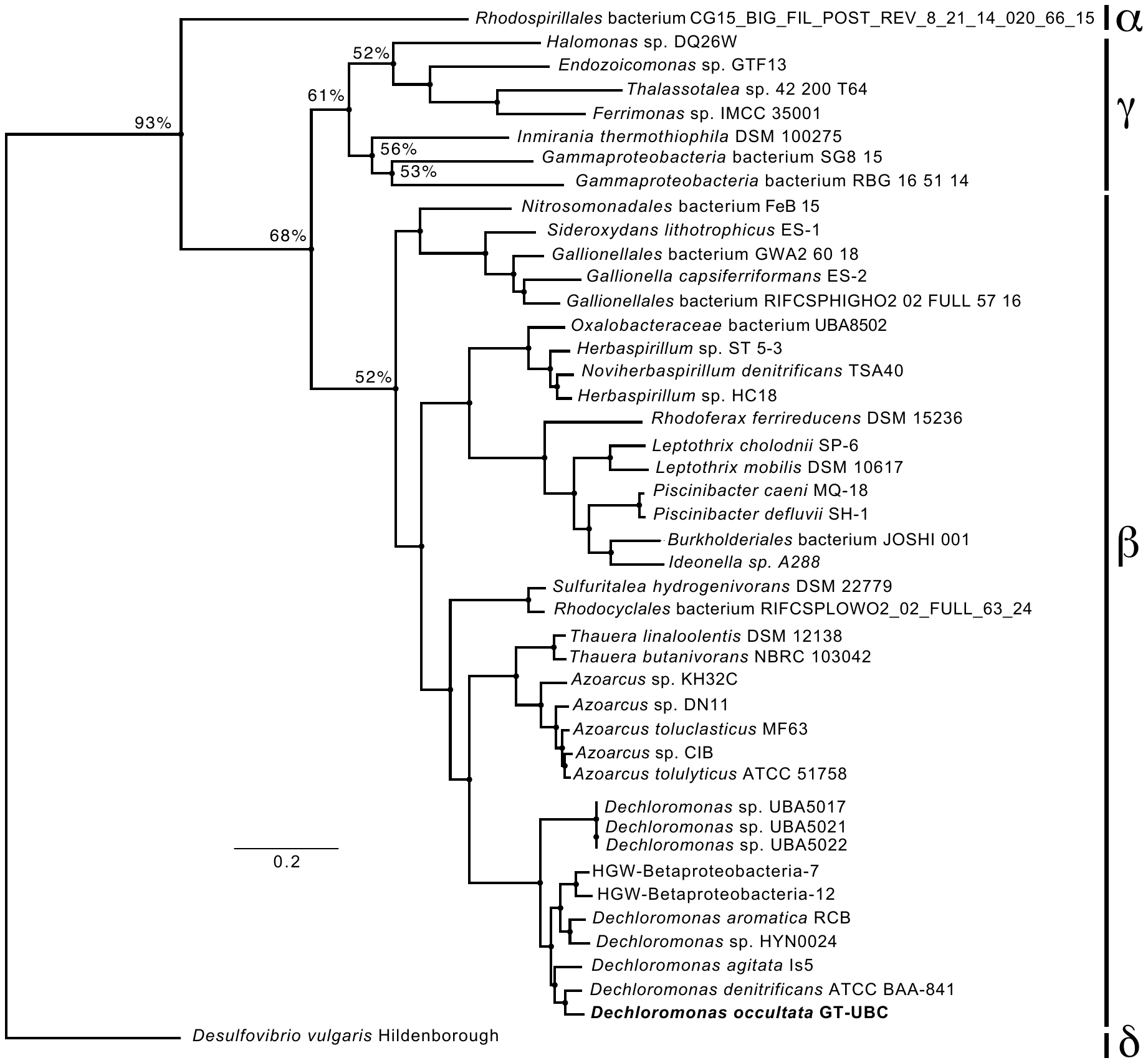

**Figure S3. Phylogeny of “*Candidatus* Dechloromonas occultata” MAG.** The phylogenic placement of the “*Ca*. D. occultata” MAG was compared to genomes of *Alphaproteobacteria*, *Betaproteobacteria* and *Gammaproteobacteria*, with *occP* and/or *mtoA* homologs **(Table S3)**. Environmental MAGs for uncultivated species are labeled with IDs. Genomes without *occP* on the phylogeny included *Dechloromonas denitrificans* (GCA_001551835.1), *Dechloromonas agitata* is5 (GCA_000519045.1), *Dechloromonas* UBA 5017 (GCA_002396525.1), *Dechloromonas* UBA 5021 (GCA_002396725.1), and *Dechloromonas* UBA 5022 (GCA_002396465.1). The deltaproteobacterium *Desulfovibrio vulgaris* was used as the outgroup (GCA_000195755.1). Bootstrap values over 50 are shown. GenBank assembly accession numbers are given in **Table S3**.

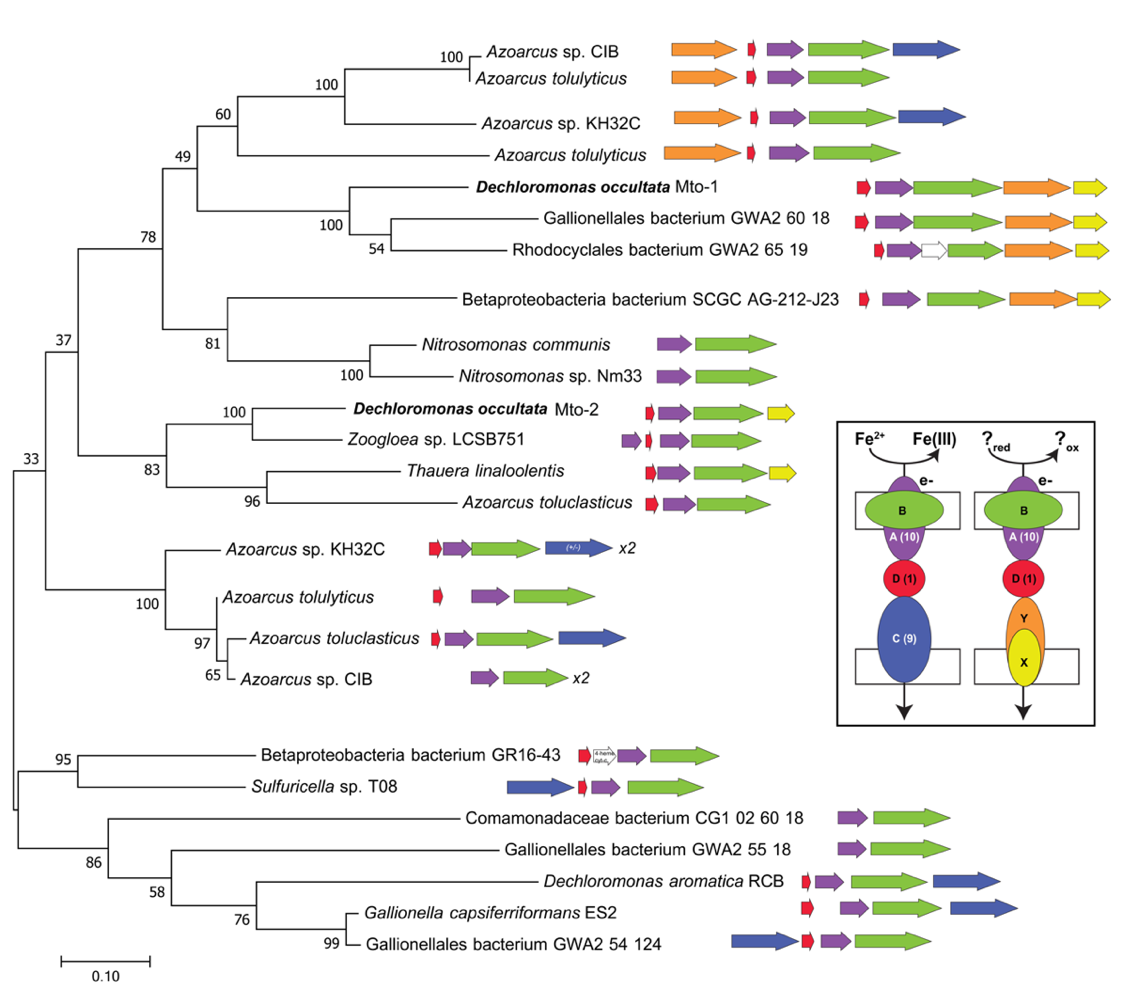

**Figure S4. Phylogeny of decaheme c-type cytochrome MtoA and synteny of Mto loci**. Maximum likelihood phylogeny of the MtoA protein sequence from “*Ca*. D. occultata” in relationship to other MtoA homologs from *Beta-* and *Gammaproteobacteria*. Accession numbers are given in **Table S3**. Bootstrap support is based on 500 samples. Next to each branch is the genomic organization of *mtoA* and neighboring genes in each species, color-coded to represent function and predicted cellular locations. Species with duplicated clusters are annotated as “x2”. Inset: left, canonical Mto pathway; right: proposed alternative Mto pathway in “*Ca*. D. occultata” and other uncultured *Betaproteobacteria*; the labels A, B, C, D, X, and Y correspond to MtoA, MtoB, MtoC, MtoX and MtoY, respectively, with heme counts in parentheses. OM: outer membrane; IM: inner membrane.

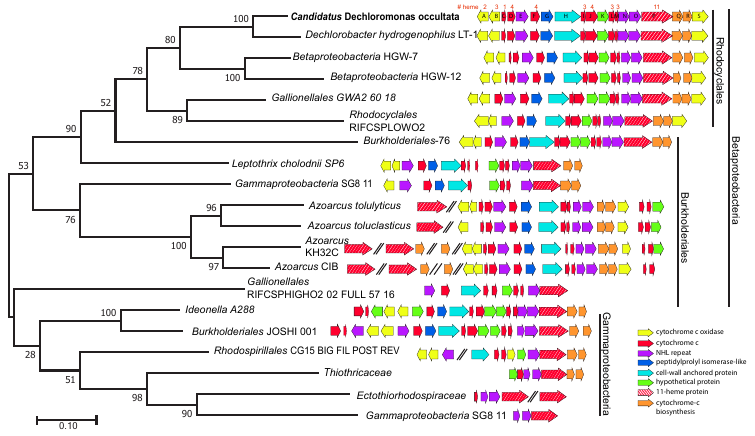

**Figure S5. Phylogeny of undecaheme c-type cytochrome OccP and synteny of Occ loci**. The tree represents the evolutionary history of the OccP protein from “*Ca*. D. occultata” in relationship to other OccP homologs from Beta- and *Gammaproteobacteria*. Accession numbers are given in **Table S3**. Note that *Gammaproteobacteria* bacterium SG8-11 contains multiple copies of the *occ* operon, one of which is within the *Burkholderiales* clade. Branch lengths represent substitutions per site. Next to each branch is the genomic organization of OccP and neighboring genes in each strain, color-coded by function.

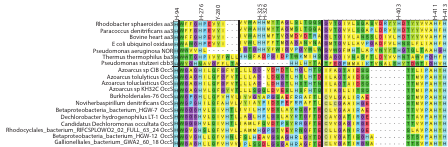

**Figure S6. Shared features of the OccS conserved domain in selected *Betaproteobacteria*.** Alignment of conserved cofactor-binding (heme a, a_3_, and Cu_B_) histidines and select D and K proton channel ligands for members of the cytochrome c oxidase subunit I family compared to OccS. Numbering is for *Paracoccus denitrificans*.

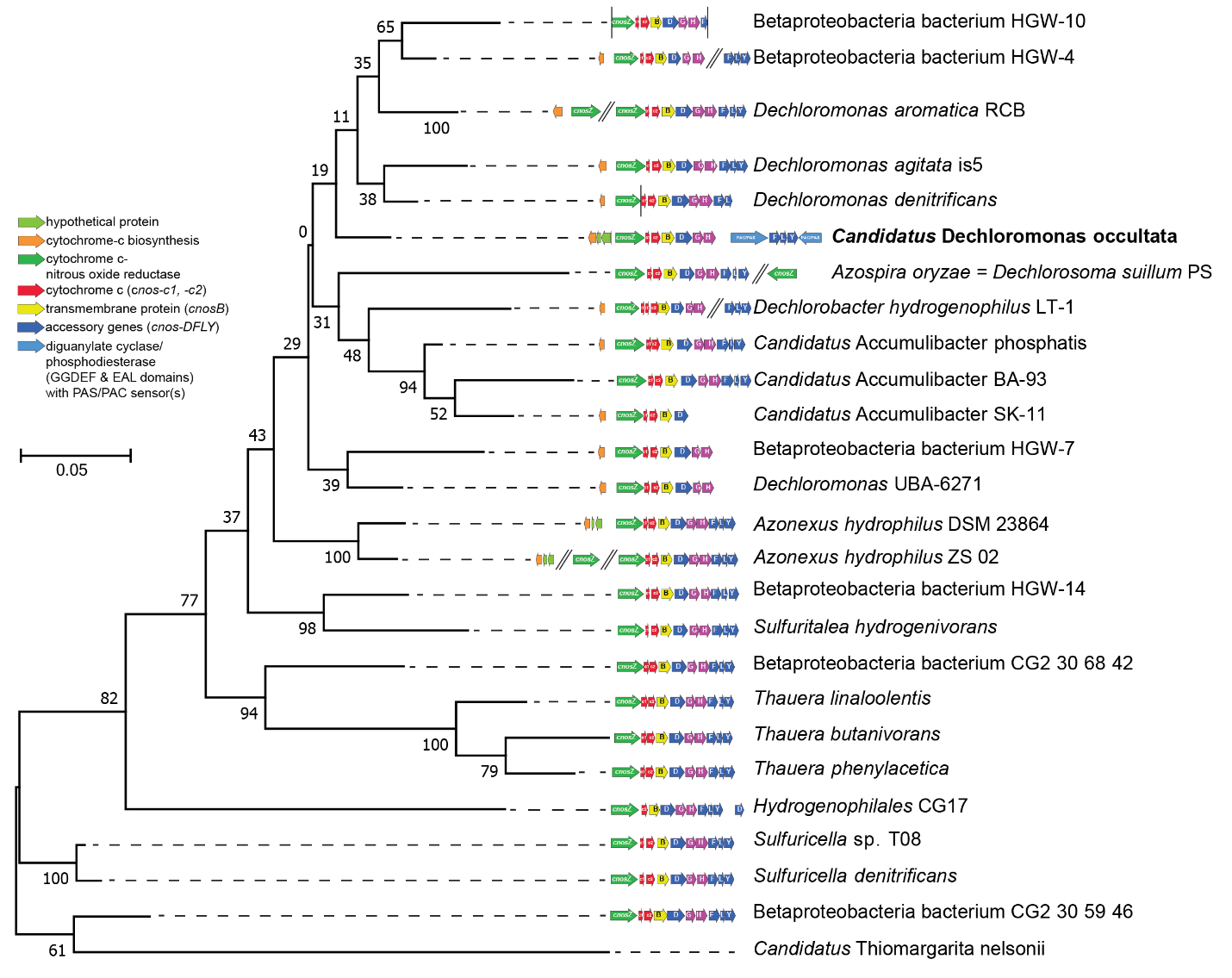

**Figure S7. Phylogeny of cytochrome-nitrous oxide reductase (cNosZ) genes and synteny of the synteny of cNosZ loci**. The tree represents the evolutionary history of the cNosZ protein from “*Ca*. D. occultata” in relationship to other cNosZ homologs from *Beta*- and *Gammaproteobacteria*. Accession numbers are given in Table 2. Branch lengths represent substitutions per site. Next to each branch we show the genomic organization of cNosZ and neighboring genes in each strain, color-coded by function.

**Figure S8.** **Abiotic reactions between Mn(III) and NH_4_^+^**. Concentrations of NH_4_^+^ (circles) and N_2_O (triangles) from 0.2 mM NH_4_^+^ added to abiotic treatments with (1 mM; closed symbols) or without (open symbols) added Mn(III) pyrophosphate. Error bars represent standard error where n=3 (N_2_O) or n=2 (NH_4_^+^).

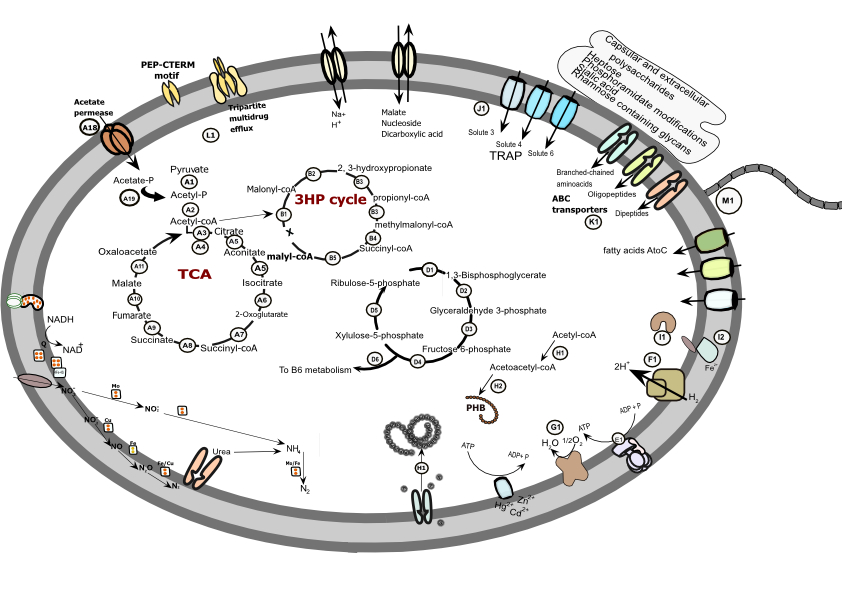

**Figure S9. “*Candidatus* Dechloromonas occultata” genomic potential and gene expression during Mn(III) reduction.** Key genes involved in central and secondary metabolism including carbon and nitrogen metabolism, energy generation, and environmental sensing are shown. Numbers correspond to proteins compiled in table S4.

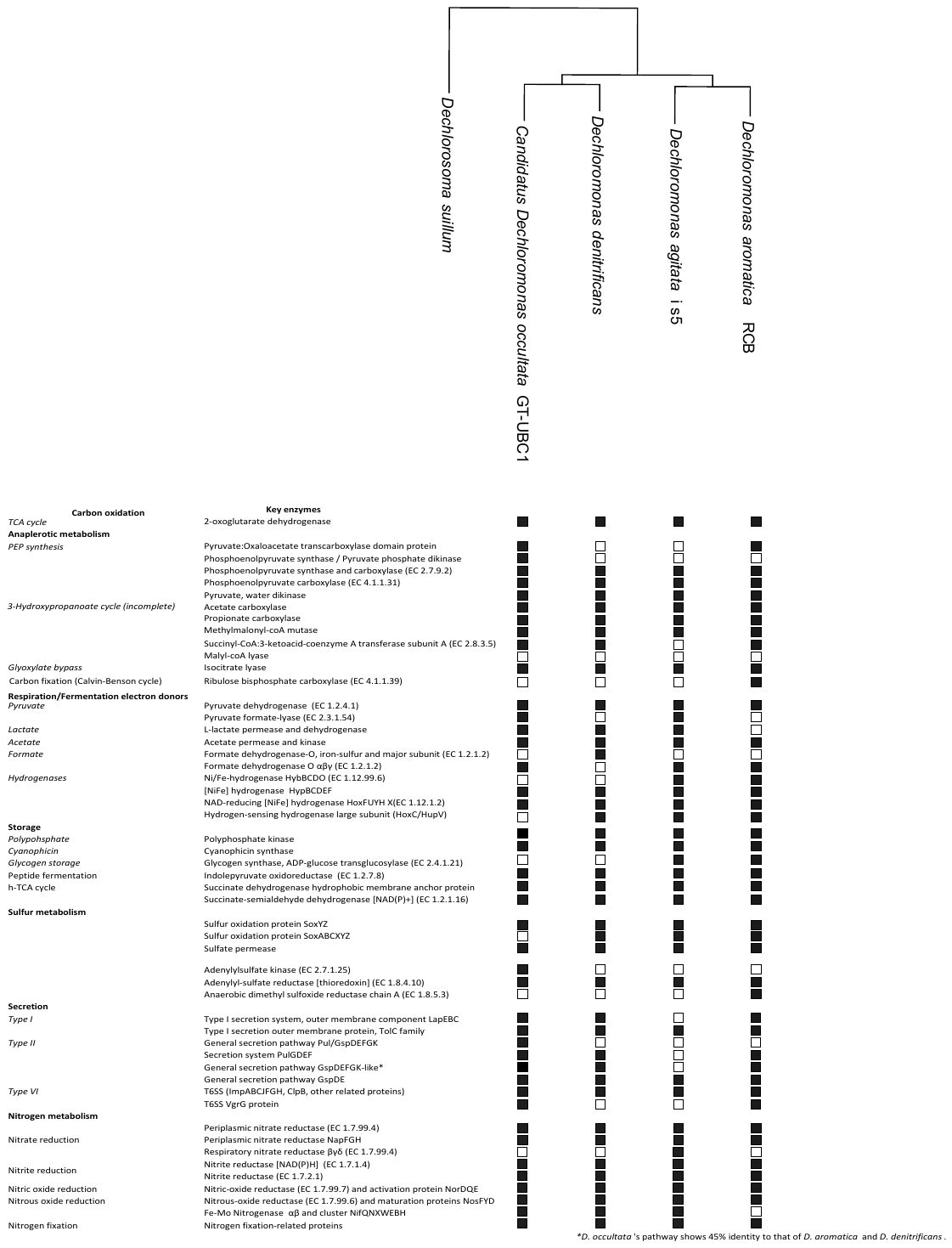

**Figure S10.** **Comparison of carbon, nitrogen and respiratory metabolic pathways for the *Dechloromonas* genus based on three representative strains and “*Ca.* D. occultata”.**

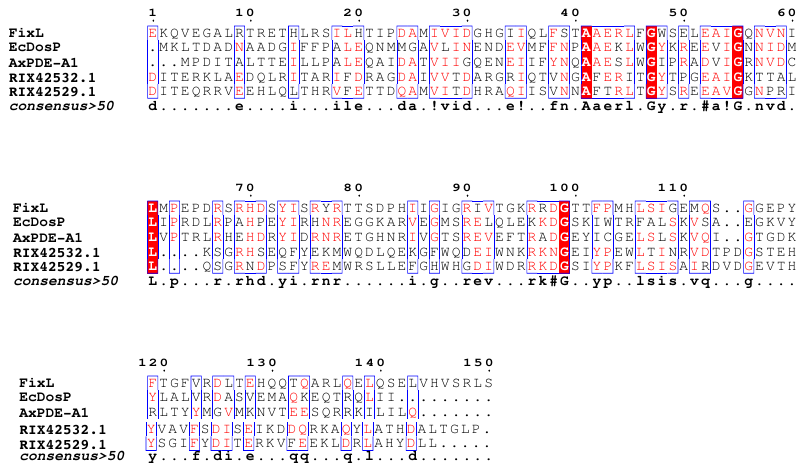

**Figure S11. Conserved features of sensor RIX42529 and betaproteobacterial homologs.** Alignment of conserved heme-binding histidines for members of the PAS-O_2_ sensor family (FixL, EcDosP, and AxPDE-A1) and two predicted PAS sensors in “*Ca*. D. occultata” (RIX42532 and RIX42529).
